# Seasonal occurrence of diamondback moths *Plutella xylostella* and their parasitoid wasps *Cotesia vestalis* in greenhouses and their surrounding areas

**DOI:** 10.1101/357814

**Authors:** Junichiro Abe, Masayoshi Uefune, Kinuyo Yoneya, Kaori Shiojiri, Junji Takabayashi

**Affiliations:** National Agricultural Research Center for Western Region, Ayabe, Kyoto, 623-0035, Japan; Center for Ecological Research, Kyoto University, Otsu, Shiga, 520-2113, Japan; Entomological Laboratory, Faculty of Agriculture, Kinki University, 3327-204, Nakamachi, Nara 631-8505, Japan; Department of Agriculture, Ryukoku University, 1-5 Ooe, Otsu, Shiga 520-2194, Japan; Department Agrobiological Resources, Faculty of Agriculture, Meijo University, Nagoya, Aichi 468-8502, Japan

**Keywords:** *Plutella xylostella*, *Cotesia vestalis*, plant volatiles, attractants, seasonal variation

## Abstract

We observed the seasonal occurrence of diamondback moth (DBM) larvae, *Plutella xylostella* (Lepidoptera: Plutellidae), and their native parasitoid wasps, *Cotesia vestalis* (Hymenoptera: Braconidae), on mizuna plants, *Brassica rapa* var. *laciniifolia* (Brassicales: Brassicaceae), in three commercial greenhouses and on wild cruciferous weeds, *Rorippa indica* (Brassicales: Brassicaceae), in the surrounding area in the Miyama countryside in Kyoto, Japan. The occurrences of DBM larvae in greenhouses followed their occurrence in the surrounding area: however, some occurrences of DBM in greenhouses took place when the DBM population in the surrounding was rather low. This suggests that the occurrence of DBM in greenhouses cannot always be explained by its seasonal occurrence in the surrounding areas. The occurrence of *C. vestalis* followed that of DBM larvae in mizuna greenhouses and in the surrounding areas. No *C. vestalis* were recorded in greenhouses when DBM was not present. *Cotesia vestalis* females preferred volatiles emitted from DBM-infested mizuna plants to those from uninfested conspecifics under laboratory conditions. Natural HIPVs (herbivory-induced plant volatiles) emitted from DBM-infested mizuna plants in greenhouses probably attracted *C. vestalis* from the surrounding area to cause their co-occurrence.

## Introduction

The rural “satoyama” forest and village landscape in Japan consists of areas of small-scale wet rice paddy fields, crop fields, and greenhouses (Kobori et al. 2003). Similar agricultural landscapes are also found in other countries (Takeuchi et al. 2003). One of the ecological characteristics of satoyama environments is that several pest insects are harbored in the surrounding natural areas (Katoh et al. 2009), and invasions of these pest insects in greenhouse is a crucial issues. In satoyama areas, carnivorous natural enemies of pest insects are also harbored in the surrounding natural areas (Kagawa and Maeto 2009).

A cruciferous crop called mizuna, *Brassica rapa* var. *laciniifolia* (Brassicales: Brassicaceae), was produced as one of the major crops in one of the satoyama areas, called Miyama, in the Kyoto Prefecture of Japan (35.3°N, 135.5°E). Pesticides were not routinely applied in the greenhouses where they were grown. The diamondback moth (DBM), *Plutella xylostella* (Lepidoptera: Plutellidae), is one of the most important pests of mizuna plants in greenhouses in Miyama. DBM and their larval parasitoids *Cotesia vestalis* (Hymenoptera: Braconidae) (Furlong et al. 2013; Talekar and Shelton 1993) are harbored in the surrounding areas (J. Abe, personal observation).

In response to damage caused by herbivorous arthropods, plants start emitting so-called “herbivory-induced plant volatiles (HIPVs)” that attract the carnivorous natural enemies of the currently infesting herbivores (Arimura et al. 2009; Dicke et al. 1990; Hare 2011; McCormick et al. 2012; Takabayashi and Dicke 1996, Takabayashi and Shiojiri 2018). The attraction capability of some of these HIPVs has been confirmed under field conditions (James 2003; James and Grasswitz 2005; James and Price 2004; Rodriguez-Saona et al. 2011; Uefune et al. 2012). We already reported that *C. vestalis* were attracted to volatiles emitted from various crucifer plants infested by DBM larvae under both laboratory (Shiojiri et al. 2000, 2010; Yoneya et al. 2018) and field conditions (Uefune et al. 2012).

We observed the seasonal occurrence of DBM larvae and *C. vestalis* on mizuna plants in three commercial greenhouses and on wild crucifer plants, *Rorippa indica* (Brassicales: Brassicaceae), in the surrounding areas in Miyama. Further, we tested whether mizuna plants infested by DBM larvae attracted *C. vestalis* by emitting HIPVs. Based on these data, we discuss the relationship between the seasonal occurrence of DBM larvae and *C. vestalis* in greenhouses and the surrounding areas.

## Materials and Methods

### Field observation

We used four mizuna greenhouses owned by a farmer, which were set in a “dice four” arrangement with 2–3 m distance between each. As there was an outbreak of DBM in one of the four greenhouses and the house was solarized, we did not used the data from that greenhouse. The greenhouses were surrounded by open agricultural fields, a thicket and a river. Since the growth stages in the four greenhouses differed, we treated the data from each greenhouse independently.

We observed the occurrence of DBM larvae and *C. vestalis* cocoons on mizuna plants in the greenhouses in 2004. Observations were made approximately every 7–14 days during the observation period. In the greenhouses, if plants had fewer than 10 leaves, then we assessed 100 plants; if they had 11–30 leaves, then we assessed 50; and for plants with more than 30 leaves, we assessed 20 plants. We also counted the numbers of DBM larvae and *C. vestalis* cocoons on a wild cruciferous species, *Rorippa indica*, which was growing in the area surrounding the greenhouses (>3 m). DBM larvae found on mizuna plants and *R. indica* plants were reared in a climate-controlled room in the laboratory (25 ± 2 °C, 50–60% RH, 16L:8D) to check the incidence of parasitism.

### Laboratory experiments

#### Insects and plants

DBM larvae were collected from fields in Ayabe, Kyoto, Japan (35°N, 135°E) in 2001, and were reared with potted komatsuna plants, *Brassica rapa* var. perviridis (Brassicales: Brassicaceae), in a climate-controlled room (25 ± 3°C, 60 ±10% RH, 16L:8D). The laboratory colony of DBM was reared on potted komatsuna plants in a climate-controlled room (25 ± 3 °C, 60 ± 10% RH, 16L: 8D) to obtain eggs. Newly emerged adults of DBM were maintained in acrylic cages (35 cm × 25 cm × 30 cm high) in a climate-controlled room (25 ± 3 °C, 60 ± 10% RH, 16L: 8D). They were provided with a 50% (v/v) honey solution as food and potted komatsuna plants to ensure mating. Komatsuna plants with eggs were collected daily and hatched larvae were reared on cut komatsuna plants in small cages (25 cm × 15 cm × 10 cm high).

*Cotesia vestalis* were obtained from parasitized DBM larvae collected in in Ayabe, Kyoto, Japan. Adults of the parasitoid species were maintained separately in plastic cages (30 cm × 20 cm × 13 cm high) with a 50% (v/v) honey solution as food in a climate-controlled room (18 ± 3 °C, 60 ± 10% RH, 16L: 8D) for 3 days to ensure mating. The second stadium DBM larvae parasitized by *C. vestalis* were put in a polypropylene box (25 cm × 15 cm × 10 cm high) with detached komatsuna leaves for food; the leaves were replaced by fresh ones every other day until the egression of *C. vestalis* larvae from DBM larvae. After egression, *C. vestalis* formed cocoons in the polypropylene box. Cocoons were collected and kept in closed-ended glass tubes until emergence. To ensure mating, emerged females were kept together with males in a plastic cage for 3 days. Thereafter, they were maintained in glass tubes (25 mm inner diameter, 120 mm length) at 18 °C to prolong their lifespan, and in continuous darkness to suppress flight. They were a maximum of 10 days old since emergence from the host. They were acclimatized for 1–2 h in the climate room before the experiments were started.

Mizuna (*Brassica rapa* var. *nipposinica* ‘Jounan-Sensuji’) and komatsuna (*B. rapa* var. *perviridis* L. ‘Rakuten’) plants were cultivated in a greenhouse (25 ± 3 °C, 60 ± 10% RH, 16L:8D). Four plants were cultivated in a plastic pot (diameter: 72 mm, depth: 65 mm) for 4–5 weeks. These potted plants were used as the odor sources in the laboratory experiments.

#### Response of C. vestalis to DBM-infested plants

Prior to each experiment, the potted mizuna or komatsuna plants had either remained uninfested or had received damage from one second stadium DBM larva per plant which had been allowed to feed for 24 hours to produce plants with at least one infested edge/leaf. Prior to the tests, the larvae, their silk, and their feces were removed from the infested plants with the aid of a fine brush.

*Cotesia vestalis* females were tested for their flight responses towards a pot of DBM-infested mizuna plants versus a pot of uninfested mizuna plants in an acrylic cage (25 × 30 × 35 cm; 3 nylon gauze-covered windows and one door) under fluorescent light (20 W, 3000 lux) in a climate-controlled room (25 ± 3 °C, 60 ± 10% RH, 16L: 8D). There was no wind in the cage. The results of Shiojiri et al. (2000, 2010) showed that visual cues are not involved in the flight responses of *C. vestalis* in the choice chamber.

Females were released individually from a glass tube (25 mm inner diameter, 120 mm length) positioned halfway between the two plant pots. Upon their first visit to one of the tested plants (defined as landing), they were removed with an insect aspirator. Ten wasps were tested using the same set of two potted plants. Each wasp was only tested once and the experiments were repeated on three or four experimental days with new sets of parasitoids and plants. We also compared the flight response of the parasitoid wasps toward a pot of DBM-infested mizuna plants versus a pot of DBM-infested komatsuna plants. Two-choice data under laboratory conditions were analyzed using the replicated *G*-test (Sokal and Rohlf 1995); parasitoids that made no choice for either plant were discarded from this analysis.

## Results

### Field observations

In the surrounding areas, *R. indica* plants were found throughout the observation period (Figure 1A: bars). The seasonal variation in the occurrence of DBM larvae in the surroundings showed two major peaks (May 9: 246 plants, 0.13 DBM/plant; June 6: 243 plants, 0.20 DBM/plant) (Figure 1A: circles). Seasonal variation in the occurrence of *C. vestalis* was also observed, with two peaks that were almost synchronous with those of the DBM (Figure 1A: squares).

**Fig. 1.**
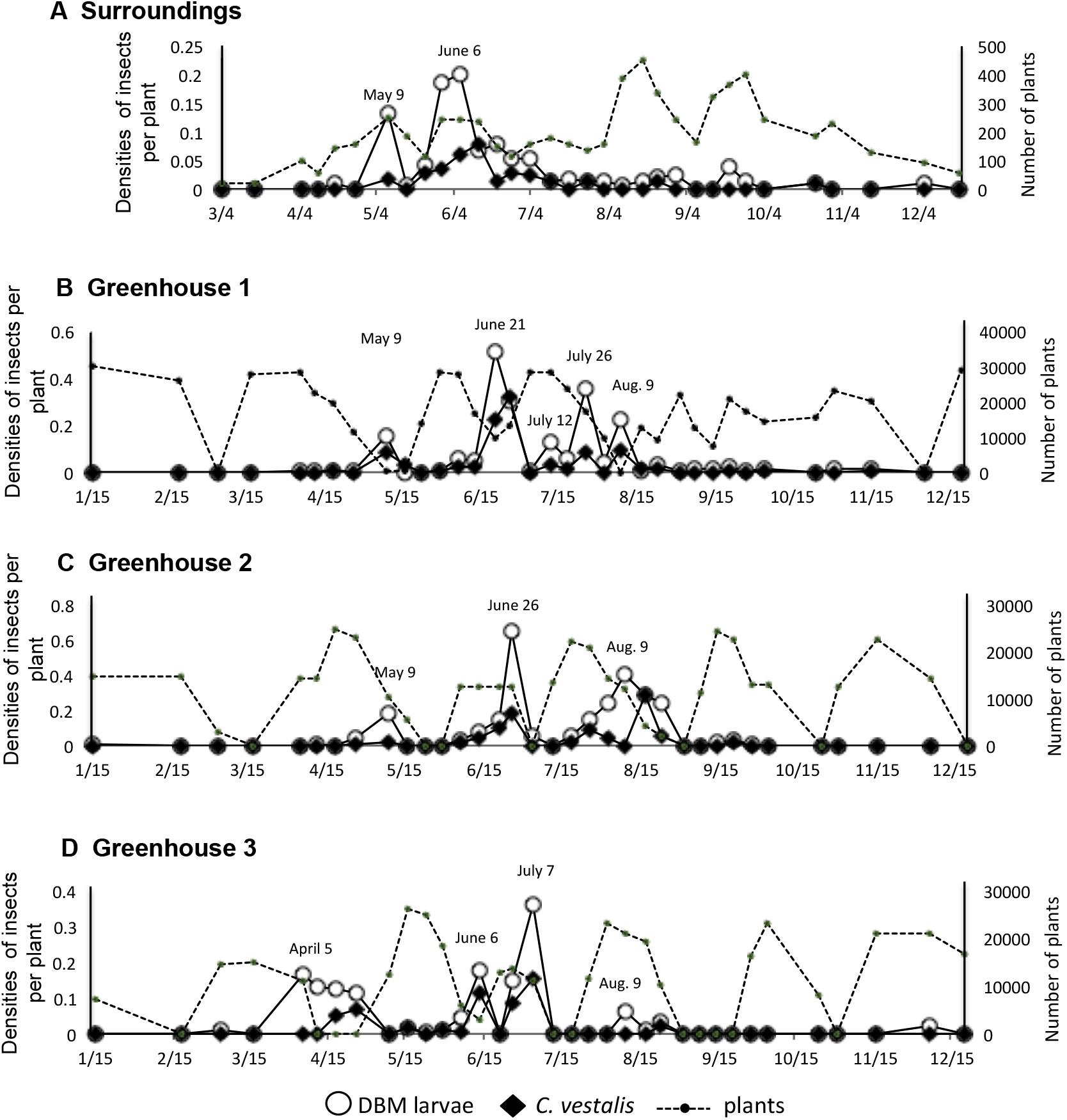
Seasonal changes to the occurrence of diamondback moth larvae (open circles) and its parasitoid wasp *Cotesia vestalis* (closed squares) in greenhouses and their surrounding areas. The numbers of host plants (mizuna and *R. indica*) (dashed lines) are also shown.

In the mizuna greenhouses (Figure 1B-E), changes in the numbers of mizuna plants (dotted lines) in the four greenhouses represented the planting and harvesting of the mizuna plants. The control threshold (level of infestation at which pesticides were applied to the crop) of DBM in a mizuna greenhouse was set at 0.05 DBM/plant (after Abe et al. 2007). Hereafter, we define the occurrence of more than 0.05 DBM/plant as a DBM peak. In the three greenhouses (Figure 1B-E), most of the plants were mizuna, with a few *R. indica* plants found solely around the edges of the greenhouses. Thus DBM peaks in the absence of mizuna plants were from *R. indica* plants in the greenhouses.

In greenhouse 1 (Figure 1B), there were five DBM peaks. The dominant peak was on June 21(9451 plants; 0.51 DBM/plant). Two peaks (July 26 and August 9) were observed when the number of DBM larvae was low in the surroundings (Figure 1A). At each DBM peak, the occurrence of *C. vestalis* was also detected.

In greenhouse 2 (Figure 1C), there were three DBM peaks. The dominant peak was on June 26 (12331 plants; 0.64 DBM/plant). One peak (August 9) was observed when the number of DBM larvae was low in the surroundings (Figure 1A). *C. vestalis* was detected at each peak.

In greenhouse 3 (Figure 1D), there were four DBM peaks. The dominant peak was on July 7 (10962 plants; 0.36 DBM/plant). Two peaks (April 5 and August 9) were observed when the number of DBM larvae was low in the surroundings (Figure 1A). *C. vestalis* was detected at each peak.

### Olfactory responses of *C. vestalis* to DBM larvae-infested plants

We offered DBM-infested mizuna plants and uninfested mizuna plants to *C. vestalis* females in a choice chamber. *C. vestalis* females preferred infested mizuna plants over uninfested mizuna plants (*G*_*P*_ = 7.1034, df = 1, *P* = 0.0077; *G*_*H*_ = 0.5977, df = 2, *P* = 0.7417; *G*_*T*_ = 7.7011, df = 3, *P* = 0.0526, replicated *G*-test) (Figure 2: upper bar). We then offered *C. vestalis* females DBM-infested mizuna plants versus DBM-infested komatsuna plants in a choice chamber. *C. vestalis* females showed an equal distribution between the two odor sources (*G*_*P*_ = 1.6108, df = 1, *P* = 0.2044; *G*_*H*_ = 0.8402, df = 2, *P* = 0.8398; *G*_*T*_ = 2.4511, df = 3, *P* = 0.6534, replicated *G*-test) (Figure 2: lower bar).

**Fig. 2.**
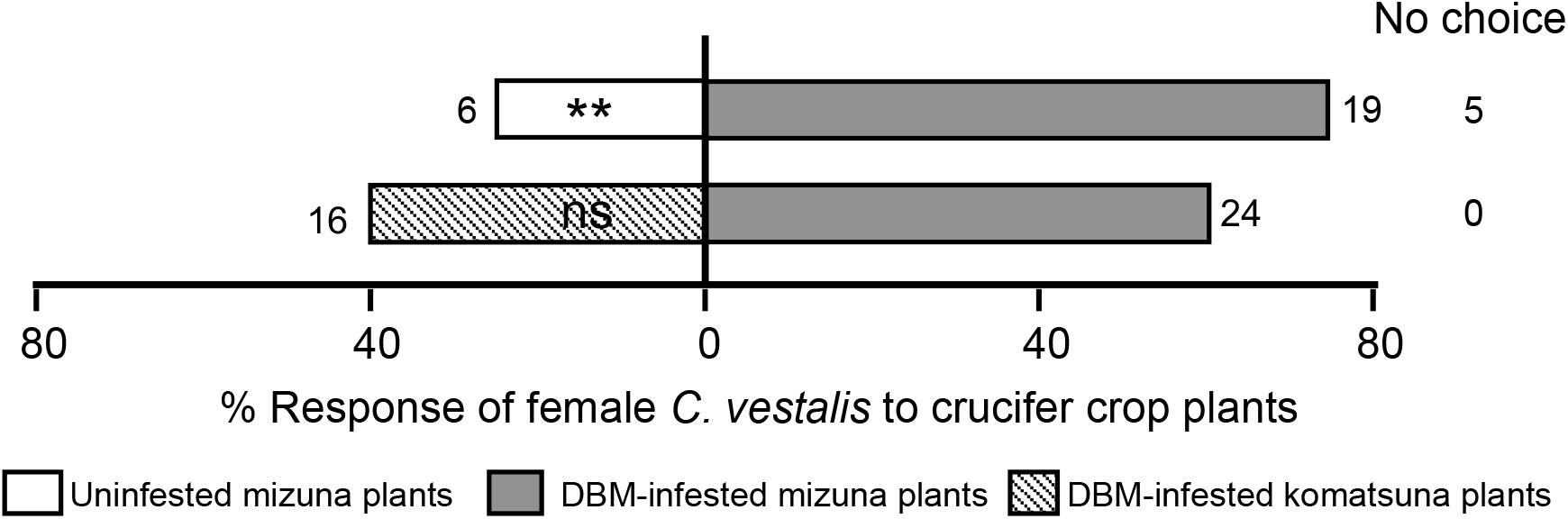
Olfactory responses of *C. vestalis* females to cruciferous plants with different treatments. The experiment was repeated on 3 experimental days (upper bar) and 4 experimental days l(ower bar); the data were pooled and subjected to a G-test. Numbers next to bars indicate the numbers of *C. vestalis* that responded to the volatiles. When the wasps did not land on either of the plants, they are called no choice individuals. ** 0.01 > P > 0.001; ns: not significantly different

## Discussion

The major DBM peaks in the greenhouses were after the DBM peaks in the surrounding areas. Interestingly, however, some DBM peaks in the three greenhouses took place when the DBM population in the surrounding areas was low. This suggested that the occurrence of DBM in greenhouses could not always be explained by the seasonal occurrence of DBM in the surrounding areas.

Throughout the observation period, DBM larvae were followed by *C. vestalis* on both mizuna plants and *R. indica* plants. This synchronized occurrence was observed in all three greenhouses when DBM occurred at a density of more than 0.05 DBM/plant. Similar synchronized occurrence is reported by Shimomoto (2002): the invasion of five native parasitoid species was observed in eggplant greenhouses in which leafminers (*Liriomyza trifolii*) occur on eggplants. *Cotesia vestalis* are attracted to volatiles emitted from various crucifer plants infested by DBM larvae under both laboratory (Shiojiri et al. 2000, 2010; Yoneya et al. 2018) and field conditions (Uefune et al. 2012). In the present study, we showed that mizuna plants started attracting *C. vestalis* after being damaged by DBM larvae, and that the attractiveness was equal to that of DBM-infested komatsuna plants (the same species as mizuna). Under experimental field conditions, Uefune et al. (2012) reported that *C. vestalis* females were attracted to HIPVs emitted from DBM-infested komatsuna plants. Taken together, volatiles from DBM-infested mizuna plants that are attractive to *C. vestalis* would be one of the factors causing the synchronized occurrence of DBM larvae and *C. vestalis* in greenhouses.

In the present study, the occurrences of *C. vestalis* in the three greenhouses were always one-step behind the occurrences of DBMs, causing a time lag between the reproduction of DBM and the removal of DBM by *C. vestalis.* Such a time lag would make *C. vestalis* less effective natural enemies in greenhouses. The continuous recruitment of *C. vestalis* into greenhouses from the surrounding areas would be one of the methods to compensate for this time lag. Four volatile compounds [(*Z*)-3-hexenyl acetate, *α*-pinene, sabinene, and *n*-heptanal] emitted from DBM-infested cabbage plants attract *C. vestalis* under both experimental greenhouse and field conditions (Ohara et al. 2017; 2018; Uefune et al. 2012). Whether the use of the four volatile compounds in greenhouses compensates for the time lag is the subject of a subsequent study.

## Author Contribution

JA, MU and JT conceived research. A, MU, YK and KS conducted experiments. JA and MU analyzed field data and conducted statistical analyses. JT wrote the manuscript. JT secured funding. All authors read and approved the manuscript.

## Acknowledgements

We thank the members of the project funded by Bio-oriented Technology Research Advancement Institution for their helpful discussions. This study was supported in part by the Bio-oriented Technology Research Advancement Institution from the Ministry of Agriculture, Forestry and Fisheries, and by grants for the scientific research of a priority area (S) and scientific research (A) from the Ministry of Education, Culture, Sports, Science and Technology of Japan (MEXT). We thank Elizabeth Kelly, MSc, from Edanz Group (www.edanzediting.com/ac) for editing a draft of this manuscript.

## References

Abe J, Urano S, Nagasaka K, Takabayashi J. (2007) Release ratio of *Cotesia vestalis* to diamondback moth larvae (*Plutella xylostella*) that suppresses the population growth of diamondback moth feeding *Brassica* leaf vegetables in a greenhouse. Bullet Nat Agr Res Centr West Region. 6:125–132. (In Japanese.)

Arimura G, Matsui K, Takabayashi J. (2009) Chemical and molecular ecology of herbivore-induced plant volatiles: Proximate factors and their ultimate functions. Plant Cell Physiol 50:911–923, doi:10.1093/pcp/pcp030

Dicke M, Sabelis MW, Takabayashi J, Bruin J, Posthumus MA. (1990) Plant strategies of manipulating predator prey interactions through allelochemicals: Prospects for application in pest control. J Chem Ecol 16(11):3091–118. doi:10.1007/BF00979614

Furlong MJ, Wright DJ, Dosdall LM, (2013) Diamondback moth ecology and management: problems, progress, and prospects. Annu Rev Entomol 58: 517–541, doi:10.1146/annurev-ento-120811-153605

Hare, JD (2011) Ecological role of volatiles produced by plants in response to damage by herbivorous insects. Annu Rev Entomol 56: 161–80. doi:10.1146/annurev-ento-120709-144753

James DG (2003) Field evaluation of herbivore-induced plant volatiles as attractants for beneficial insects: Methyl salicylate and the green lacewing, *Chrysopa nigricornis*. J Chem Ecol 29: 1601–1609, https://doi.org/10.1023/A:1024270713493

James, DG and Grasswitz, T R (2005). Field attraction of parasitic wasps, *Metaphycus* sp. and *Anagrus* spp. using synthetic herbivore-induced plant volatiles. BioControl 50: 871–880, DOI 10.1007/s10526-005-3313-3

James DG, Price TS (2004) Field-testing of methyl salicylate for recruitment and retention of beneficial insects in grapes and hops. J Chem Ecol 30: 1613–1628 doi:10.1023/B:JOEC.0000042072.18151.6f

Katoh K, Sakai S, Takahashi T (2009) Factors maintaining species diversity in satoyama, a traditional agricultural landscape of Japan. Biol Conserv 142: 1930–1936, doi.org/10.1016/j.biocon.2009.02.030

Kagawa Y, Maeto K (2009) Spatial population structure of the predatory ground beetle *Carabus yaconinus* (Coleoptera: Carabidae) in the mixed farmland-woodland satoyama landscape of Japan. Eur. J. Entomol 106: 385–391, doi:10.14411/eje.2009.049

Kobori H, Primack RB (2003) Participatory conservation approaches for satoyama, the traditional forest and agricultural landscape of Japan. Ambio. Jun;32: 307-11

McCormick AC, Unsicker SB and Gershenzon J (2012) The specificity of herbivore-induced plant volatiles in attracting herbivore enemies. Trends Plant Sci 17: 303–310

Ohara, Y., Uchida, T., Kakibuchi, K., Uefune, M. and Takabayashi, J. (2017) Effects of an artificial blend of host-infested plant volatiles on plant attractiveness to specialist parasitic wasps. J Appl Entomol 141: 231–234, doi:10.1111/jen.12328

Ozawa R., Ohara, Y., Shiojiri K., Uchida, T., Kakibuchi, K., Kugimiya S., Uefune, M. and Takabayashi, J. (2018) Uninfested plants and honey enhance the attractiveness of a volatile blend to a parasitoid *Cotesia vestalis*. J Appl Entomol (in press)

Rodriguez-Saona C, Kaplan I, Braasch J, Chinnasamy D, Williams L (2011) Field responses of predaceous arthropods to methyl salicylate: a meta-analysis and case study in cranberries. Biol Control 59:294–303 doi.org/10.1016/j.biocontrol.2011.06.017

Shimomoto, M. (2002) Control of *Liriomyza trifolli* (Burgess) by indigenous and imported parasitoids on eggplant in forcing culture. Bull. Kochi Agric, Res. Cent. 11: 37–44

Shiojiri, K, Takabayashi, J, Yano, S and Takafuji A (2000) Flight response of parasitoid toward plant-herbivore complexes: a comparative study of two parasitoid-herbivore systems on cabbage plants. Appl Entomol Zool 35: 87–92, doi.org/10.1303/aez.2000.87

Shiojiri K, Ozawa R, Kugimiya S, Uefune M, van Wijk M, Sabelis MW and Takabayashi J (2010) Herbivore-specific, density-dependent induction of plant volatiles: Honest or “cry wolf” signals? PLoS One e12161 doi.org/10.1371/journal.pone.0012161

Sokal RR and Rohlf FJ (1995) Biometry: the principles and practice of statistics in biological research, 3rd ed. W.H. Freeman and Company; New York.

Takabayashi J, Dicke M (1996) Plant-carnivore mutualism through herbivore-induced carnivore attractants. Trends Plant Sci 1:109–113, doi.org/10.1016/S1360-1385(96)90004-7

Talekar NS, Shelton AM, 1993. Biology, ecology, and management of the diamondback moth. Ann Rev Entomol 38, 275–301.

Takeuchi, K., Brown, R. D., Washitani, I., Tsunekawa, A. and Yokohari, M. (2003) Satoyama: The traditional rural landscape of Japan. Springer-Verlag

Uefune M, Choh Y, Abe J, Shiojiri K, Sano K and Takabayashi J (2012) Application of synthetic herbivore-induced plant volatiles causes increased parasitism of herbivores in the field. J Appl Entomol 136: 561–567, doi:10.1111/j.1439-0418.2011.01687.x

Yoneya K, Uefune M, Takabayashi J (2018) Parasitoid wasps’ exposure to host⍰infested plant volatiles affects their olfactory cognition of host⍰infested plants. Anim Cogn 21:79–86, doi:10.1007/s10071-017-1141-3

